# TDP-43 α-helical structure tunes liquid-liquid phase separation and function

**DOI:** 10.1101/640615

**Authors:** Alexander E. Conicella, Gregory L. Dignon, Gül H. Zerze, Hermann Broder Schmidt, Alexandra M. D’Ordine, Young C. Kim, Rajat Rohatgi, Yuna M. Ayala, Jeetain Mittal, Nicolas L. Fawzi

## Abstract

Liquid-liquid phase separation (LLPS) is involved in the formation of membraneless organelles (MLOs) associated with RNA processing. Present in several MLOs, TDP-43 undergoes LLPS and is linked to the pathogenesis of amyotrophic lateral sclerosis (ALS). While some disease variants of TDP-43 disrupt self-interaction and function, here we show that designed single mutations can enhance TDP-43 assembly and function via modulating helical structure. Using molecular simulation and NMR spectroscopy, we observe large structural changes in a dimeric TDP-43. Two conserved glycine residues (G335 and G338) are potent inhibitors of helical extension and helix-helix interaction, which are removed in part by variants including the ALS-associated G335D. Substitution to helix-enhancing alanine at either of these positions dramatically enhances phase separation *in vitro* and decreases fluidity of phase separated TDP-43 reporter compartments in cells. Furthermore, G335A increases TDP-43 splicing function in a mini-gene assay. Therefore, TDP-43 helical region serves as a short but uniquely tunable module that shows promise as for controlling assembly and function in cellular and synthetic biology applications of LLPS.

## Introduction

RNA-binding proteins harboring low-complexity sequences are major constituents of membraneless organelles, particularly ribonucleoprotein (RNP) granules^1–3^. These granules can display liquid-like properties^4^, consistent with the view that they assemble through liquid-liquid phase separation (LLPS)^5^. The molecular interactions and physical forces by which RNP granule proteins assemble into liquid-like compartments has been of great interest^6–10^. Multivalent interactions based on several interaction motifs can give rise to LLPS, including globular domains (often in modular chains) binding linear protein or nucleic acid motifs, as well as weak multivalent interactions between polar, aromatic, and charged-patterned disordered protein segments^11,12^. The diversity of interaction types mediating LLPS could in principle allow for different responses to stimuli, as well as variations in permeability, dynamics, and persistence, thus allowing comparable diversity in their respective biological functions^13–15^.

TDP-43, a well-known RNP granule component^16–19^, is a heterogeneous ribonucleoprotein (hnRNP) family member with genetic and histopathological links to amyotrophic lateral sclerosis (ALS)^20^, frontotemporal dementia (FTD)^21^, and Alzheimer’s disease (AD)^22^. The tandem RRM domains recognize UG-rich sequences found near splice sites^23^, contributing to TDP-43’s role in regulating splicing of transcripts in mammalian cells^24,25^. Yet, the splicing function of TDP-43 also depends on the TDP-43 globular N-terminal domain^26–30^ and its predominantly intrinsically disordered C-terminal domain (CTD, residues 267-414)^31^. One may wonder what is the role of these domains? TDP-43 CTD, rich in polar and aromatic residues, is prone to self-assembly and confers TDP-43 the ability to undergo LLPS both *in vitro*^32^ and in cells^33^. Though LLPS requires the polar CTD, LLPS and function of TDP-43 is further enhanced by intermolecular interactions mediated by the folded N-terminal domain and by an evolutionarily conserved partially *α*-helical subregion of the CTD spanning residues 320-343. This region is required for LLPS of TDP-43 reporter constructs^33^ and important for splicing in a well-established mini gene assay^31^. The importance of the conserved region to TDP-43 function is underscored by the presence of 7 ALS-associated variants in this short stretch^34^.

Previously we showed that the conserved TDP-43 CTD region mediates intermolecular *α*-helical contacts enhancing TDP-43 phase separation^32^. In particular, a transiently helical region spanning 321-330^35^ is stabilized and extends into the adjacent 331-343 region upon self-association^32^. Importantly, we found that several ALS-associated missense mutations in this region (including Q331K and M337V) disrupt helix-helix interaction, impairing phase separation^32^. Intriguingly, the adjacent 331-343 region that becomes partially helical upon assembly contains two nearby conserved glycine positions, G335 and G338. Because inclusion of designed helical regions have been recently demonstrated to enhance protein phase separation^36^, we suspected that these “helix-discouraging” residues may serve as important sites regulating TDP-43 assembly. Therefore we set out to test if TDP-43 helical structure and helix-helix interaction can be tuned to enhance phase separation and increase TDP-43 function by α-helix enhancing mutations at conserved glycine positions.

In this work, we use experiments and simulations to characterize the influence of these two nearby glycine positions on TDP-43 CTD, structure, self-association, phase separation *in vitro* and in cells, and splicing function. We also test the effect of the G335D ALS-associated mutation occurring at one of these sites to understand a potential mechanism leading to disease^37^. Here we pair an extensive experimental and computational structural and biophysical characterization of TDP-43 CTD including a number of mutants targeted at these two glycine residues with an evaluation of the role of these positions on TDP-43 LLPS and splicing function in cells. The combination of high-resolution NMR spectroscopic and computational structural data, phase separation approaches *in vitro* and in cells, and functional data provide an unprecedented and comprehensive atomically detailed picture of the tunability of phase separation, connecting fine control of the sequence, to structure, to interactions and finally to both phase separation and function.

## Results

### Helical structure of TDP-43 CTD enhanced by dimerization

Our previous work showed that the conserved region (CR) in TDP-43 CTD (residues 320-343) encompasses a region from 321-330 that is partially helical and that helicity is enhanced in 321-330 and spreads to 331-343 upon TDP-43 self-interaction^32^. Yet, the precise extent of increased α-helicity and structural change in the assembled state remains poorly understood, despite the importance of the CR in TDP-43 splicing function^31^ and phase separation^32,33^. Because we could not trap the assembled state, we previously relied on experiments sensitive to the small population of assembled state (chemical shift deviation vs. concentration, and relaxation dispersion NMR) present in equilibrium with the monomeric state. Signs of enhanced helicity were present, but direct characterization of the size of, and extent of helicity in, the assembled form could not be determined^32^. Here, to directly visualize the structural changes along the self-assembly pathway, we first used all-atom explicit solvent simulations. Using advanced sampling techniques (see Methods), we determined the equilibrium ensemble of configurations of two copies of a computationally-tractable segment of TDP-43 CTD, TDP-43_310-350_, containing the entire helical region that we previously characterized and validated^32^ placed together in a simulation box. First, we found that overall *α*-helical content for the entire two-chain ensemble (i.e. including configurations with and without intermolecular contacts) is greater (up to 60% helical from residues 321-331, Figure S1) than that of the single chain ensemble (up to 50% helical) from our previous work. Next, to search for stable configurations populated in the two chain ensemble, we computed a free energy surface quantifying the relative energetic favorability of different configurations binned along two order parameters selected to represent intermolecular assembly: the number of hydrogen bonds and number of hydrophobic contacts between the two protein chains (Figure 1A). Two distinct minima (ΔG values *<*10 kJ/mol relative to most populated reference bin), which we term strongly and weakly bound states, are observed. Averaging over only the configurations corresponding to the strongly bound free energy well (i.e. configurations in favorable intermolecular contact), we see a larger increase in *α*-helical content (Figure 1B) and stabilization (~25% population) of a continuous helix spanning from residues 320-334, abruptly ending at residue G335 (Figure 1C). The simulations suggest heterogeneity in the bound ensemble. Many different residue pairs in the region spanning 320-343 form intermolecular contacts (Figure S1A). The strongly bound ensemble also comprises heterogeneous binding orientations including parallel and antiparallel helix-helix interactions as well as interactions involving disordered conformations of the 320-343 region (Figure S1B,C). Though two chains may not be sufficient to explore the full ensemble of bound configurations (i.e TDP-43 may form trimeric or higher order helical assemblies), these data demonstrate how self-interaction can promote helical stabilization and are consistent with our previous NMR paramagnetic relaxation enhancement data which suggested that TDP-43 CTD 320-343 region visits multiple bound orientations^32^. In summary, stabilization of helical elements upon binding suggests a self-assembly mechanism involving both “conformational selection” and “induced fit” as proposed previously for other helix-based assemblies^38^.

**Figure 1:**
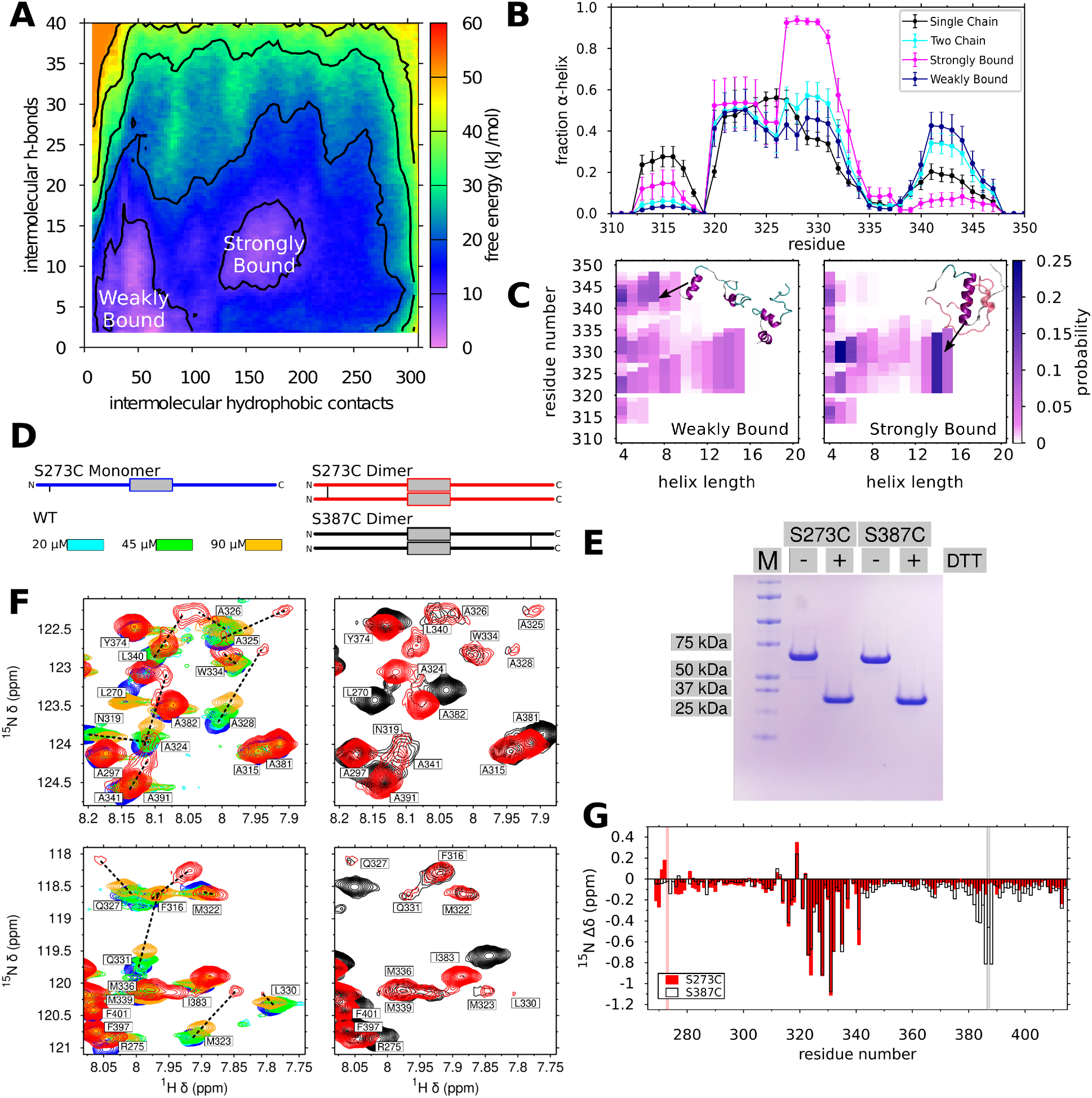
TDP-43 CTD self-associates and forms transient helical structures. A) Free energy landscape of TDP-43_310-350_ intermolecular contacts for all-atom two-chain simulations highlight two minima corresponding to strongly and weakly bound states. B) α-helical content of TDP-43 simulations at each residue. Data are plotted as mean ± s.e.m of n=5 equal divisions of the total data set. C) *α*-helical secondary structure propensity maps show the contribution of all helical configurations to the overall helical content shown in B. This analysis highlights stabilization of contiguous *α*-helical structure in the strongly-bound state. D) Schematic of crosslinked CTD variants. E) Non-reducing SDS-PAGE of dimeric S273C and S387C illustrate an apparent reduction of molecular weight by approximately one half upon addition of disulfide-breaking reducing agent (1 mM DTT). F) (Left) Select regions from ^1^H-^15^N TROSY spectra of dilsulfide-linked homodimeric TDP-43 CTD S273C (red) show line broadening and large upfield ^15^N chemical shift differences in 321-343 region compared to monomeric reduced S273C (blue), consistent with formation of structure. Chemical shifts for residues in the dimer state were assigned by extrapolation (dashed black line) from spectra of 20 *μ*M (cyan), 45 *μ*M (green), and 90 *μ*M (orange) WT. (Right) Overlay of ^1^H-^15^N TROSY spectra regions for homodimeric S273C and S387C are similar in the central 320-340 region. G) Quantification of ^15^N Δ*δ* values between homodimeric S273C (red) and S387C (black) and their respective monomers. Gray boxes indicate the positions of S273C and S387C disulfide crosslinks.

Our previous work demonstrated that even in the absence of phase separation, the conserved region of TDP-43 CTD self-assembles mediated by helix-helix contacts into a multimer state invisible to standard NMR approaches^32^. To experimentally trap a dimer state, we instead created a novel approach, mimicking the two chain simulations by forming obligate CTD dimers *in vitro* by covalently crosslinking two CTDs far (>40 residues) from the conserved 320-343 region (Figure 1C). We created two distinct disulfide cross-linked dimers, oxidized by copper (II) phenanthroline, at single cysteines engineered at residues 273 or 387 (Figure 1C). After oxidation, these CTD variants appear at twice the apparent molecular weight compared to reduced control samples in non-reducing SDS-PAGE (Figure 1D). Consistent with stabilization of a structured assembly, fingerprint two dimensional (^1^H-^15^N TROSY) spectra of both S273C dimers (10 μM, in monomer units) and S387C dimers (20 μM) reveal shifted, very weak resonances that can be assigned to the 320-343 region by extrapolation of resonances of the WT TDP-43 CTD as a function of increasing concentration (i.e. increasing population of an assembled state) (Figure 1E). This observation strongly suggests that the cross-link does not bias the helix-helix contacts into a particular geometry (i.e. parallel assembly due to symmetric cross-link site), possible due to the long (40 residue) region between the cross-link site and the helical region. Quantification of ^15^N chemical shift differences between dimeric (oxidized) and monomeric (reduced) CTD revealed upfield chemical shifts from residues 315 to 343, primarily in 321-335, as well as at the site of the crosslink (Figure 1F). Importantly, both cysteine variants display the same pattern of upfield shifts in the 321-340 region (RMSD ~ 0.07 ppm), indicating that the chemical environment and hence the dimeric states sampled by the 321-340 region are independent of crosslink position. Furthermore, the magnitude of the ^15^N chemical shift differences observed here are similar to those we derived for the assembled state of TDP-43 CTD based on experimental CPMG relaxation dispersion NMR experiments^32^. Interestingly, these obligate dimer samples were not stable at higher concentrations, precluding more extensive NMR characterization. Taken together, the computer simulations and experiments show signatures of extensive enhanced helicity upon dimerization focused in the region from residue 321 to 335.

### Mutations at glycine positions enhance CTD *α*-helical structure in the 330-343 region

Given that TDP-43 conserved region helicity seems to be disrupted in the region of G335 and G338, we wondered if we replaced G335 or G338 with other residues, would we increase helicity? To test the hypothesis that substitution of residues that are less helix-discouraging would enhance TDP-43 CTD helicity, we designed several TDP-43 CTD variants at G335 and G338 positions to observe the effect of these amino acids on TDP-43 CTD helicity. First, we conducted single chain all-atom molecular dynamics simulations of the same 41-residue fragment, TDP-43_310-350_, with a set of single point mutations, changing G335 or G338 from glycine to the helix-promoting residue alanine (G335A and G338A). Mutating glycine to alanine (G335A and G338A) results in a significant increase of *α*-helical content in the simulations, particularly in the vicinity of the mutation sites (Figure 2A). The increase in helical content is highest within residues 331-343 (Figure 2B, right and Figure S2A,B), while helical content in residues in the preceding partially helical region 321-330 does not increase (Figure 2B, left). G335D, an ALS-associated variant, has similarly enhanced helicity from 331-343, as do engineered variants G335N (similar to G335D but with the uncharged amide analogous residue) and G335S, suggesting that the enhancements in helicity caused by G335A are due to the removal of the unique backbone conformational flexibility of glycine, not specific side-chain mediated contacts. Quantifying the helical enhancement, the maximum length of contiguous *α*-helical conformations^39^ increases for G335A (i.e. helices spanning 320-337) and the population of longer helical conformations increases (Figure 2C). For G338A, long helices are populated in the region including position 338 that are not observed for WT.

**Figure 2:**
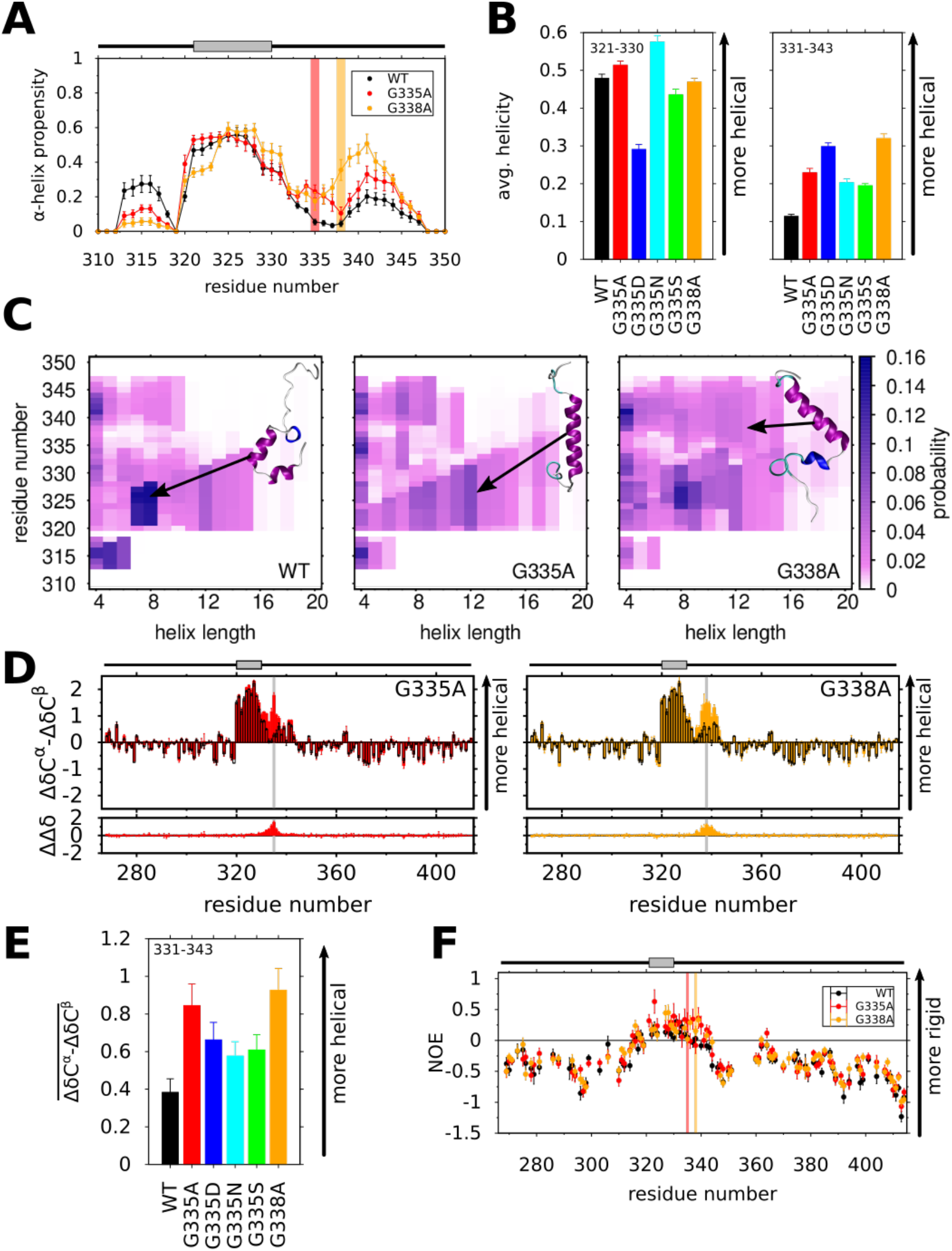
G335 and G338 substitutions enhance in TDP-43 CTD helical region enhance helical stability. A) Per-residue *α*-helical propensities for WT, G335A, and G338A for atomistic simulations of TDP-43_310-350_ show higher fraction of *α*-helical structure near the sites of mutation (highlighted by red, G335A, or orange, G338A, bars). Data are plotted as mean ± s.e.m of n=5 equal divisions of the total data set. B) Average simulated helicity is mostly unaffected for residues 321-330 (left) away from G335 and G338 mutation sites but enhanced for 331-343 (right) surrounding the mutation site. C) Maps of contiguous *α*-helical structure location (y-axis) and length (x-axis) for TDP-43_310-350_ show increased probability for longer helix structure in G335A and G338A relative to WT. One example configuration from a subpopulation of the simulation ensemble (indicated by the black arrow) is displayed for each variant. D) Experimental NMR secondary chemical shifts (**ΔδC**^***α***^ − **ΔδC**^***β***^) and the differences in secondary shifts with respect to WT (ΔΔ*δ*) for G335A and G338A show increased *α*-helical structure near the site of mutation. The secondary shift values for WT are overlaid in black for comparison. E) The average experimental secondary shift 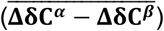 for TDP-43 331-343 of WT and mutant CTD highlight increases in local *α*-helical structure in all variants. Error bars are SEM. F) Higher {^1^H}-^15^N heteronuclear nuclear Overhauser effect (NOE) values measured for G335A (red) and G338A (orange) compared to WT (black) near the site of mutation indicate slowed local protein backbone motion, consistent with enhanced helical structure. Mutation positions G335A and G338A highlighted with gray bars. For all above panels: Simulation data are plotted as mean and SEM from 10 equal blocks of equilibrated structural ensemble. Unless otherwise stated, experimental data are plotted as mean and SD estimated from signal to noise ratio.

With strong computational support for enhanced helicity by G335 and G338 point variants, we next characterized the secondary structure of TDP-43 CTD variants experimentally by comparing NMR chemical shifts of WT and mutant TDP-43 CTD variants. Positive values of the difference of the residual ^13^Cα and ^13^Cβ chemical shifts compared to a random coil reference database, ΔδC^*α*^ − ΔδC^*β*^, (Figure 2D) indicate population of α-helical structure. Relative to WT, all of the G335 and G338 point mutants show positive increases in secondary shifts, ΔΔδ (Figure 2D and Figure S2C) consistent with the enhanced helical content predicted by simulation. As suggested by the simulations, the pattern of local *α*-helical enhancement is distinct for G335A and G338A; enhancements are observed from 330-339 for G335A and from 335-343 for G338A (Figure 2D). Importantly, the values of ΔδC^*α*^ − ΔδC^*β*^ near the site of mutation nearly reach the level seen in the 321-330 region, which is approximately 50% helical^32^. To compare the helical enhancement between all variants, we computed the average secondary shift across residues 331-343, 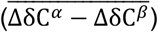 (Figure 2E). Average secondary shift values for residues 331-343 of WT and mutant CTD are strongly correlated (r = −0.972) with the predicted change in free energy of helix stabilization relative to glycine^40^ (Figure S2D). To further probe for enhanced helical structure in TDP-43 CTD, we measured NMR relaxation parameters sensitive to reorientational motion at each position, which are slower for structured conformations than for disordered conformations. Sensitive to motions on the <1 ns timescale, heteronuclear nuclear Overhauser effect values, {^1^H}-^15^N NOE, for G335A and G338A are slightly higher than WT in the region from 330-343 (Figure 2F), consistent with slower motions due to higher population of structured, slow moving conformations. Similarly, the value of the transverse relaxation rate constant, ^15^N *R*_2_, is higher for G335A and G338A, consistent with slower motions. Because ^15^N *R*_2_ is also enhanced by transient helix-helix contacts in TDP-43 (Figure S2E, see below) we also measured the cross-correlated relaxation rate, *η*_xy_, which is independent of these exchange effects. Like the {1H}-^15^N NOE, *η*_xy_ is enhanced from 331-343 in G335A and G338A, consistent with slower motion in the 331-343 region. Differences in ^15^N spin relaxation parameters for WT, G335A, G338A are negligible across the remainder of the CTD, suggesting that the variants do not change overall folding of the monomeric CTD (e.g. they do not promote the formation of new tertiary structure such as intramolecular helix bundling). Taken together, these data strongly suggest that single point variants at conserved glycine positions in TDP-43 331-343 enhance TDP-43 CTD helicity with a magnitude commensurate with their predicted ability to stabilize α-helix structure.

### G335 and G338 mutations enhance intermolecular helix-helix contacts and higher-order assembly

In our previous work, we showed that ALS-associated variants Q331K, M337V, A321G, and A321V disrupt intermolecular helix-helix assembly of TDP-43 CTD^32^. Here, we wondered if TDP-43 CTD variants that enhance helicity in 331-343 region increase TDP-43 CTD assembly. For this purpose, we used the same approach, measuring the concentration-dependent perturbations of NMR resonances in fingerprint (^1^H-^15^N HSQC) spectra for G335 and G338 variants at concentrations ranging from 10 *μ*M to 90 *μ*M at conditions where phase separation does not occur (i.e. 0 mM NaCl). We then calculated chemical shift perturbation (Δ*δ*) at each concentration relative to a monomeric reference for each variant (i.e. the concentration below which we do not detect significant chemical shift differences = 20 *μ*M for all variants except for G338A = 10 *μ*M; Figure 3A). As shown previously^32^, WT CTD displays upfield ^15^NΔ*δ* in the 321-340 region above 20 *μ*M, indicative of local intermolecular interactions. G335A and G338A show ^15^N Δ*δ* at the same residue positions as WT but are significantly enhanced at 30 *μ*M compared to WT. Above 30 *μ*M, many of the resonances in G335A and G338A are broadened beyond detection (Figure 3A), consistent with enhanced intermolecular interactions, compared to WT which shows resonance loss due to broadening only above 90 *μ*M.

**Figure 3:**
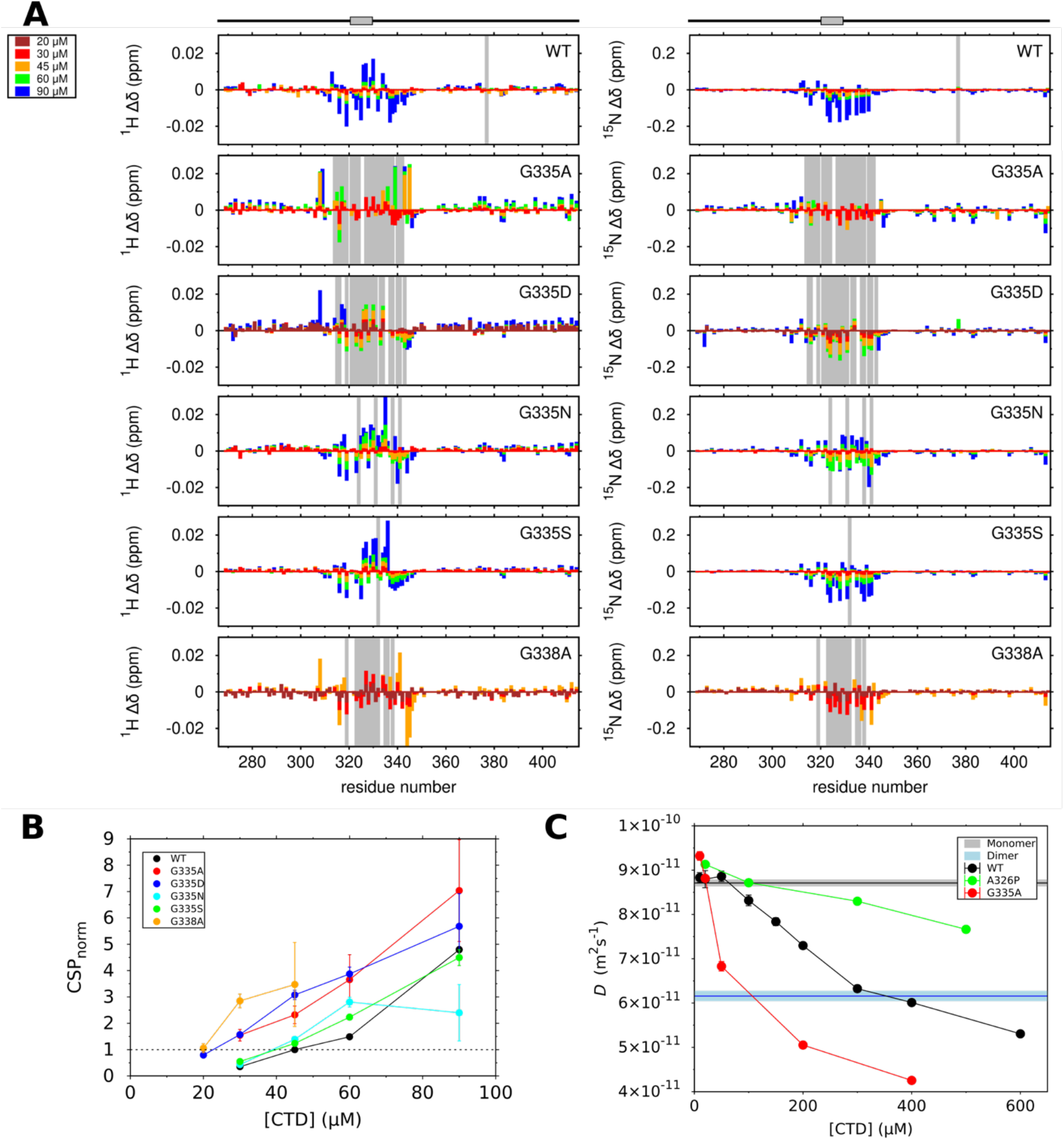
Mutations enhance TDP-43 CTD helix-helix interaction and assembly. A) Larger ^15^N chemical shift differences (Δ*δ*) in the helical region of TDP-43 CTD for G335A and G338A compared to WT are consistent with enhanced helix-helix interactions. Shifts are reported at concentrations from 20 *μ*M to 90 *μ*M with respect to a monomeric (low concentration) reference (20 *μ*M for WT and G335A; 10 *μ*M for G338A). Gray boxes indicate the loss of detectable peak intensity due to line broadening. B) Normalized chemical shift differences vs concentration are higher for mutant CTD compared to WT, consistent with increased self-assembly for G335 and G338 variants. Error bars are SD estimated by 100 bootstrapping simulations derived from experimental uncertainty. C) Diffusion coefficients calculated from NMR diffusion data plotted as a function of CTD concentration show that WT (black) and G335A (red) diffuse slower than A326P (green). Average diffusion coefficients for S273C and S387C dimers and S273C and S387C monomers are indicated by blue and black horizontal lines, respectively. Data are plotted as mean ± s.d., indicated for monomer and dimer diffusion coefficients by gray and light blue boxes, respectively.

To quantify the effects of G335 and G338 variants on intermolecular self-assembly in the 321-343 region, we used linear regression to normalize the relative magnitude of ^15^N Δ*δ* for WT and mutant CTD variants at each concentration to the ^15^N Δ*δ* observed for WT for 45 *μ*M - 20 *μ*M, which we termed CSP_norm_ (Figure 3B). CSP_norm_ is higher for all G335 and G338 variants relative to WT, suggesting that these mutations enhance intermolecular interactions in the 321-343 region. G338A displays the largest increase in CSP_norm_ values followed by G335A and G335D, and weaker enhancements caused by G335S and G335N. As we previously observed for WT^32^, CSP_norm_ values for all variants increase approximately linearly as a function of increasing CTD concentration, suggesting that the population of assembled state is not saturated at these concentrations. Importantly, the enhancement of helix-helix assembly due to each variant follows the same order as the enhancement in helicity of the monomeric CTD; the only exception is G335D whose lower net charge likely favors assembly in these low salt conditions^41^. Together these observations suggest that higher helicity of TDP-43 CTD monomer enhances helix-helix assembly by increasing the effective affinity.

High concentration samples that contain significant populations of the assembled form, however, show significant line broadening in the region of the helix for all variants, precluding our ability to probe saturation of binding and to determine the dissociation constant for assembly. Instead, we took advantage of the remaining resonances far from the assembly site and still visible at high concentration to probe the average hydrodynamic size as a function of concentration. To this end, we measured translational diffusion coefficients for WT CTD at sample concentrations ranging from 10 *μ*M to 600 *μ*M. Intensity ratios for all concentrations decay as a single exponential as a function of squared gradient strength (Figure S3), suggesting that the translational diffusion of the CTD at each concentration can be best captured by a single diffusion coefficient (*D*), reflecting the population-weighted average diffusion rates for monomeric and oligomeric states consistent with dynamic, reversible assembly^42^. At low CTD concentrations (50 *μ*M and below) the apparent diffusion coefficient for WT CTD remains largely unchanged (8.8± 0.1 · 10^−11^ m^2^s^−1^) (Figure 3C), suggesting that this value reflects the diffusion coefficient of monomeric CTD. This diffusion coefficient corresponds to an approximate radius of gyration of 2.8 nm in dilute aqueous solution, close to that of the similar-length low complexity (LC) domain of hnRNPA2^43^ and significantly more extended than the globular reference protein lysozyme (2.03 nm) (Figure S3A), which is of almost identical molecular weight to TDP-43 CTD. At concentrations above 50 μM, however, the apparent diffusion coefficient of the CTD sample decreases, consistent with CTD assembly into higher order species. These diffusion data alone, however, do not uniquely determine the oligomerization state. To provide a reference for the diffusion of a TDP-43 dimer, we also measured the translational diffusion coefficients of the monomeric and covalent dimeric forms of the S273C and S387C CTD variants at low (5 μM) concentrations (Figure 3C, Figure S3B). Diffusion coefficients are similar for both cross-linked dimers. The diffusion coefficient of both reduced monomeric CTD (8.7 ± 0.1 · 10^−11^ m^2^s^−1^) is effectively the same as the wild-type whereas the covalent dimer is smaller (6.4 ± 0.1 · 10^−11^ m^2^s^−1^). As concentrations exceed 300 *μ*M, the change in diffusion coefficient of wild-type CTD decreases, suggesting saturation of assembly. The diffusion coefficient at high concentrations is also smaller than that of the covalently linked dimer, suggesting that oligomers larger than dimers are present. For the G335A variant, oligomerization occurs at much lower concentration and the apparent diffusion coefficient also plateaus at high concentration, implying saturation of a similarly-sized state. Hence, G335A self-associates at lower concentrations, implying tighter binding. To demonstrate that higher order structures, not increased nonspecific contacts or increased viscosity due to increased protein concentration, give rise to the slower diffusion coefficients observed, we tracked the diffusion of the helix-disrupting A326P variant^32^ that does not assemble (Figure 3C). This variant shows only modestly slower diffusion even at high concentration, and the profile lacks a sigmoidal shape, suggesting that the sigmoidal shape for WT and G335A arises due to assembly in the concentration range of the transition. Taken together, these data suggest TDP-43 CTD assembles into states larger than dimers and that such an assembly happens at lower concentrations for the interaction-enhancing G335A variant.

### G335 and G338 variants alter *in vitro* LLPS

Given the relationship between TDP-43 CTD intermolecular self-assembly and LLPS, we next tested if G335 and G338 variants that increase interaction also enhance phase separation. Although the conserved region mediates helix-helix contacts at all conditions, TDP-43 CTD does not readily phase separate in the absence of salt, which appears to induce or enhance interaction between the polar-residue regions of TDP-43 as well as other prion-like domain proteins^32,43,44^. To monitor the effect of G335 and G338 variants on phase separation, we mapped the low concentration arm of the phase diagrams as a function of increasing NaCl using a recently developed approach^7,43,45^. In our case, LLPS was initiated by brief incubation of 20 *μ*M CTD with a given concentration of NaCl, followed by centrifugation to sediment the micron-sized droplets making up the phase-separated state (i.e. the protein dense phase). We then measured the concentration of protein in the supernatant (i.e the protein dispersed phase) for each condition, representative of the saturation concentration of TDP-43 CTD (i.e. the concentration of CTD above which LLPS will occur at these conditions). The amount of TDP-43 CTD remaining in the supernatant decreases for WT CTD as a function of NaCl concentration, suggesting CTD is undergoing LLPS (Figure 4A) with “salting-out” behavior, consistent with our previous work on TDP-43 CTD^32^ as well as FUS LC^44^ and hnRNPA2 LC^43^. Both the ALS-associated variant Q331K as well as the LLPS-disrupting control variant M337P display higher remaining protein in the supernatant than WT for any given salt concentration, consistent with our previous findings that these mutations discourage CTD assembly and phase separation^32^. Conversely, G335 and G338 variants all show lower concentration of protein remaining in the supernatant compared to WT, suggesting that these single point variants are significantly more prone to phase separate. Micrographs of the mutant CTD after phase separation show no morphological differences of these variants compared to WT and no evidence of increased protein aggregation (Figure 4B) suggesting that changes in protein remaining in the supernatant faithfully report on TDP-43 CTD LLPS and not on a transition to aggregation in these conditions. Of our engineered CTD variants, G335A and G338A show the greatest enhancement of LLPS, with only about 5 *μ*M remaining in the supernatant at 150 mM NaCl (i.e. physiological ionic strength) conditions, a nearly 3x decrease compared to WT (~15 *μ*M). The G335D, G335N, and G335S variants displayed more modest enhancements, with approximately 10 *μ*M remaining. Thus, single substitutions can dramatically enhance phase separation via enhancement of helix-helix contacts. Importantly, these saturation concentrations for TDP-43 CTD variants show a strong correlation with changes in CTD secondary structure (Figure 4C) more so than with amino acid hydropathy (Figure S4A,B), suggesting that *α*-helical structure stabilized by the amino acid at that position, and not hydrophobicity changes, accounts for the large changes in LLPS. Interestingly, only G335D shows a different rank-order in this saturation concentration as a function of increasing salt concentration – at low salt concentrations (<150 mM NaCl), the saturation concentration for G335D is lower (i.e. greater propensity to phase separate) than that of WT, G335N, and G335S, while at the highest salt concentrations tested (300 mM to 500 mM NaCl), it is approximately the same as WT and is higher (less prone to phase separate) than G335N and G335S. This change as a function of salt may arise because G335D decreases the net charge of TDP-43 CTD, leading to less long range repulsion at low salt conditions lacking salt to screen these effects^41^.

**Figure 4:**
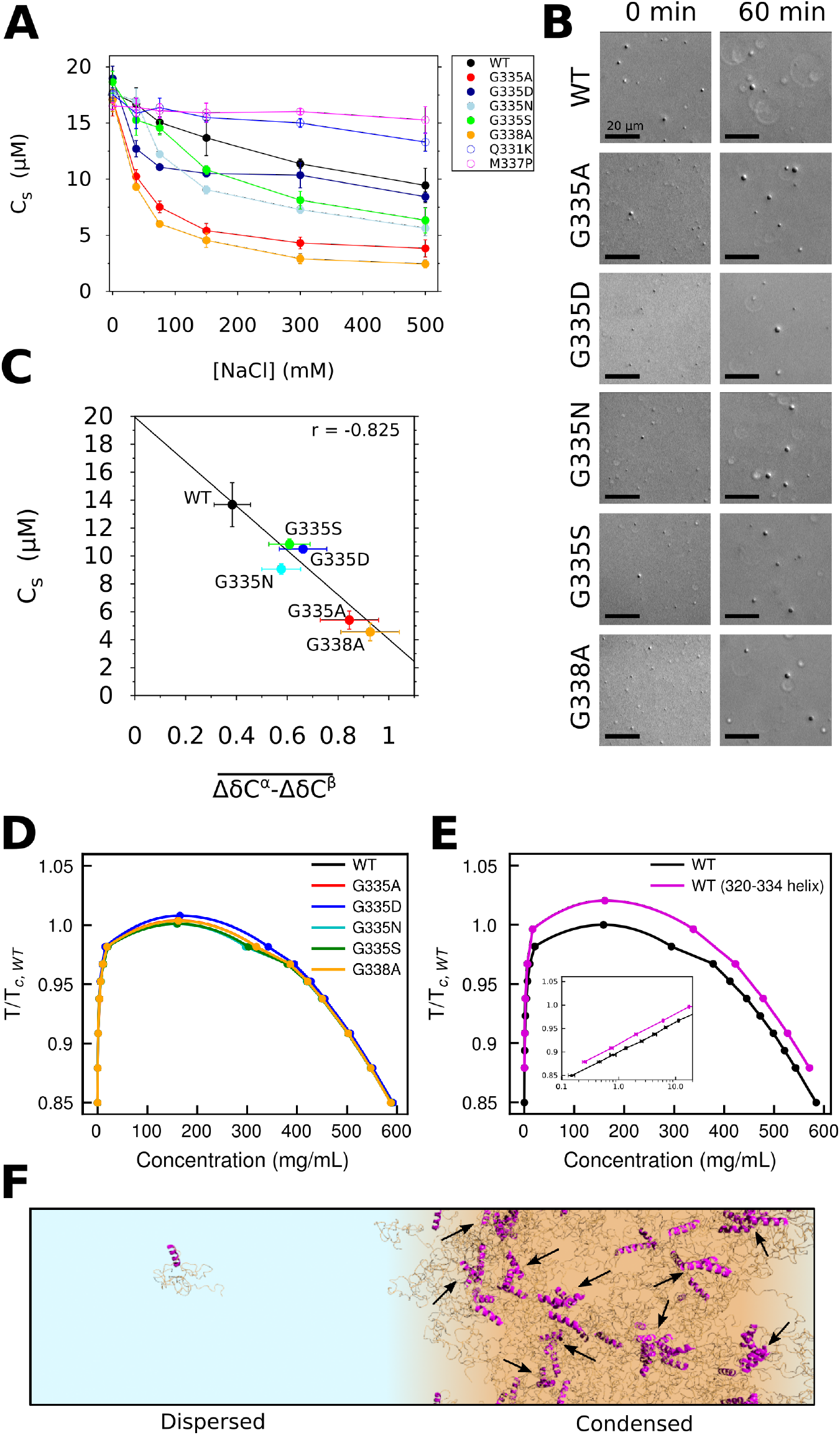
Mutations at G335 and G338 alter *in vitro* LLPS. A) Phase diagrams for WT and mutant CTD measured for 0 to 500 mM NaCl. Error bars represent SD if three replicates. B) The saturation concentration (C_s_) of WT and mutant CTD at 150 mM NaCl (“physiological conditions”) correlates well with the average change in secondary shift 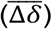. Error bars represent SD of three replicates for C_s_ and SEM for 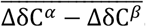. C) DIC micrographs of LLPS for WT TDP-43 CTD and variants G335A, G335D, G335N, G335S, and G338A at 0 min and 60 min after addition of NaCl to 300 mM. Scale bar is 20 *μ*m. D) Coexistence curves of TDP-43 CTD from coarse-grained (CG) slab simulations of WT and G335/G338 mutants illustrate no discernable difference in LLPS behavior as a result of mutation. E) Coexistence curves of TDP-43 from CG slab simulations illustrate reduced thermal miscibility when residues 320-334 are treated as a rigid body (magenta) instead of fully flexible (black). Inset shows the ~2-fold change to left arm of the phase diagram with temperature vs. concentration (on a log scale). F) A snapshot of the TDP-43 CTD CG slab simulation where CTD molecules harboring rigid *α*-helical substructure (magenta) visit both dispersed and condensed states in phase coexistence. Arrows indicate oligomers formed by helix-helix contacts.

To further elucidate the potential contributions of hydrophobicity vs. helicity to TDP-43 phase separation, we conducted coarse-grained simulations of TDP-43 CTD coexistence following the simulation model and framework from our previous work^46–48^. Because this coarse-grained model treats protein chains as flexible polymers and secondary structure is not directly captured, this allows us to decouple the mutational change in hydrophobicity from the changes in helicity, which are inseparable in experiment and traditional all-atom molecular simulation. Even in the absence of helicity in the conserved region, self-interaction at the 310-340 region is slightly favored as seen by elevated contact probability in coarse-grained simulations of the condensed phase (Figure S4C). Changing only the amino acid sequence in the coarse-grained model (e.g. G335A) and hence the favorability of interactions, there is no appreciable change in the phase diagram generated from directly simulating phase coexistence (Figure 4D). We then tested the effects of the *α*-helical configuration by using rigid body constraints to effectively force the subregion identified by 2-chain all atom simulation (residues 320-334; Figure 1B) into an *α*-helical configuration (Figure 4E). When we compare the results from fixed-helix coexistence simulations (Figure 4F) and those using fully flexible chains, we see that the phase diagram shifts upward to higher temperatures (Figure 4E) indicating a greater phase separation propensity for the fixed-helix chains. Upon introduction of *α*-helices, the saturation concentration, indicated by the left arm of the coexistence curve, decreases by roughly a factor of 2 across a broad range of temperatures tested (Figure 4E, inset). These results provide direct evidence that the presence of short *α*-helical structure can promote self-interaction and hence enhance phase separation of TDP-43 CTD, more so than just the increase of sidechain interaction strength.

### G335 and G338 variants affect in-cell LLPS fluidity and splicing function

To elucidate how changes in conserved region helicity impact TDP-43 phase behavior and function in a cellular context, we tested the G335 and G338 point mutations in the context of established cell-based reporter assays for TDP-43 phase dynamics and splicing function. We first introduced G335 and G338 mutations into the TDP-43_RRM-GFP_ construct, previously developed to determine the effect of single point mutations on TDP-43 phase fluidity in cells by fluorescence recovery after photobleaching (FRAP) of the single, large nuclear TDP-43 reporter droplet phase formed in each cell nucleus, into which partitions the great majority of the expressed TDP-43_RRM-GFP_^33^. We measured fluorescence signal recovery for WT and mutant droplets over time (Figure 5A-D). For comparison, we included only droplets of similar size and signal levels in our analysis. Previously tested CTD variants that disrupt helix-helix interaction show increased fluidity due to decreased intermoleuclar interaction, while removal of the conserved segment prevented phase separation^33^. Here, we now show that CTD mutants G335N, G335S, and G335A instead display decreased droplet fluidity relative to WT (Figure 5B,D). G335A shows the greatest reduction of fluidity, while G335N and G335S show intermediate droplet fluidity. Importantly, the rank order of the in cell fluidity matches that of the *in vitro* C_sat_ (see Figure 4) (except for G335D, see below), though some observations were unexpected, highlighting the importance of the cellular context. In particular, FRAP profiles were variable among G335S droplets for reasons not clear at this point (Figure S5A) and the magnitude of the difference in fluidity between G335S and G335N is not predicted from the *in vitro* TDP-43 CTD LLPS C_sat_. Regarding the disease-associated variant G335D, the in cell FRAP recovery time of G335D droplets is not significantly different from that of WT (Figure 5 B,D), similar to the *in vitro* LLPS at high salt. Taken together, these findings show that increasing CR helix propensity by mutating G335 and G338 not only lowers C_sat_ for phase separation *in vitro*, but also reduces TDP-43 fluidity in a tunable manner correlated with the properties observed *in vitro*. Our data furthermore suggest that C_sat_ and droplet material properties can be precisely tuned by single amino acid changes in the helix-helix interacting region.

**Figure 5:**
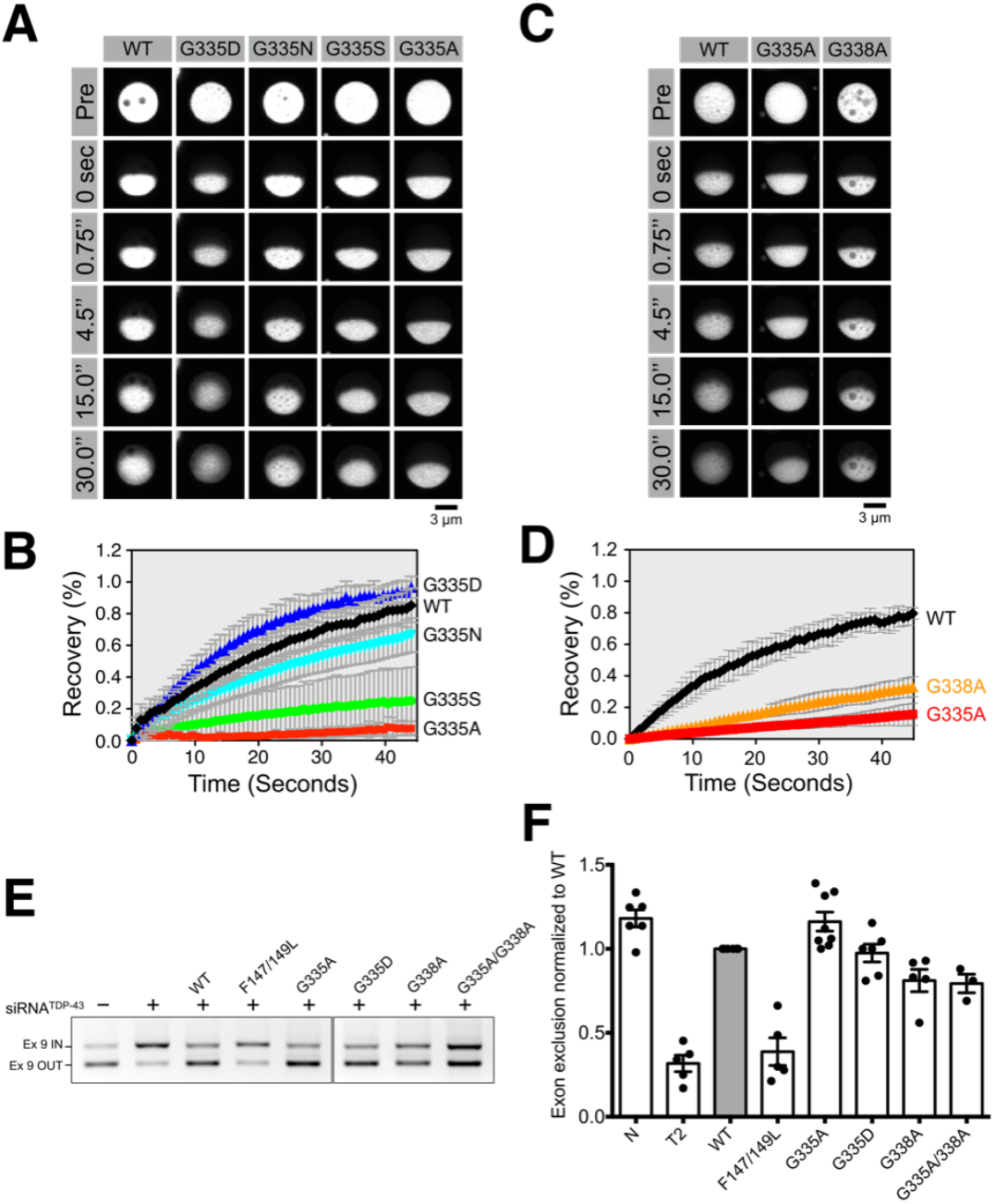
Mutations at G335 and G338 alter in-cell LLPS fluidity and splicing function. A-D) Fluorescence recovery after in cell half-droplet photobleaching and quantification for TDP-43_RRM-GFP_ reporter construct with (A,B) WT, G335D, G335N, G335S, and G335A or (C,D) WT, G335A, and G338A CTD sequences. TDP-43_RRM-GFP_ shows differential droplet fluidity for G335 variants relative to WT. E) Example agarose gel of PCR products to quantify the fraction of exon 9 CFTR exclusion (OUT) expressed from a minigene construct transfected in HeLa cells. Cells were treated with control, TDP-43 siRNA or TDP-43 siRNA plus WT or mutant vectors. F) The splicing activity was calculated as the ratio of exon 9 exclusion relative to WT. Error bars are SEM of 3 or more replicates.

**Figure 6:**
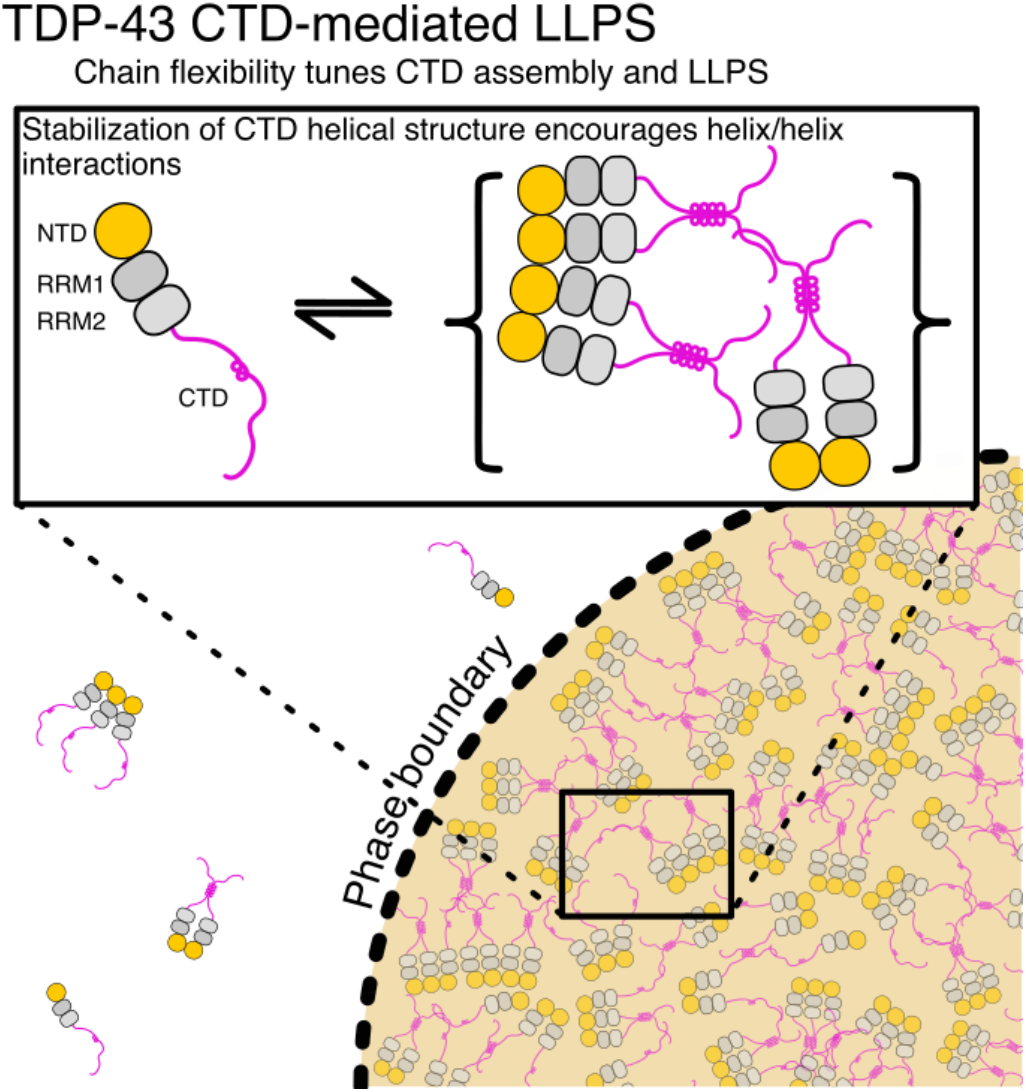
A model for contributions of CTD chain flexibility to TDP-43 LLPS. The TDP-43 CTD is sufficient for assembly into sparsely populated oligomeric states and LLPS states at low and high salt concentrations, respectively. Engineered or disease-causing mutations that stabilize CTD *α*-helical structure result in increased helix/helix intermolecular interactions and phase separation.

Considering the links between TDP-43 assembly and splicing function^26,27^, we next sought to investigate whether mutations at G335 and G338 can alter its ability to facilitate exon skipping in the cystic fibrosis transmembrane conductance regulator (CFTR) minigene splicing assay^49^. Previously it was shown that deletion of the conserved region containing the helix in TDP-43 CTD disrupted RNA splicing^31^. Here, we wanted to test if enhanced TDP-43 CTD helicity and helix-helix interaction not only promote phase separation but also alter splicing function. To this end, we performed rescue experiments in TDP-43 knock-down HeLa cells comparing the regulation of CFTR exon 9 splicing by mutant and WT TDP-43. We measured the extent of exon skipping in siRNA treated cells expressing control, siRNA-resistant WT, G335, or G338 mutants (Figure 5E, S5C). The percent of exon exclusion was normalized to WT to calculate the relative splicing regulatory activity for each mutant (Figure 5F, S5D). As a negative control, we included the RNA-binding deficient variant F147L/F149L in our assays. Importantly, G335A variant shows increased activity compared to WT (p value < 0.004), suggesting that a single point substitution can indeed enhance TDP-43 function. Contrary to our expectations and unlike the enhancement observed for G335A, G335D and G338A show slightly decreased splicing activity compared to WT. This difference between G335A and G338A in splicing effect may be due to differential alterations in interactions with binding partners including hnRNPA2 mediated via the CR of TDP-43 CTD that contribute to TDP-43 splicing activity^31^ or by non-natural over-stabilization of the TDP-43 interaction. Indeed, in cellular stress conditions, G338A (and G335A) appear to increase cytoplasmic mislocalization and aggregation of full-length TDP-43 compared to WT (Figure S5B). Thus, even though phase separation and splicing activity can be increased in cells by point mutations in the conserved region, TDP-43 self-interaction, droplet fluidity and splicing function may be precisely balanced by sequence to prevent aggregation and preserve splicing function.

## Discussion

Diverse structural elements spanning the spectrum from completely disordered regions to globular domains have been shown to contribute to the interactions that undergird biomolecular LLPS. The role of the conserved region within TDP-43 CTD comes into focus here. We show that the α-helical structure of the region can be increased in direct proportion to the α-helix stabilizing propensity of the amino acid substituted at a glycine position, with the helix-promoting alanine substitution having the greatest effect. These data suggest that the backbone conformational entropy discourages helix formation in the 331-343 region of TDP-43, as substitution of glycine 335 with any of several small amino acids (including G335A, G335S, G335N, and G335D) all enhanced helical structure. This helix stabilization in turn enhances TDP-43 CTD helix-helix interaction and phase separation. These data complement our previous report that variants in the same subsection of the conserved region spanning residues 331-343 that becomes helical only upon helix-helix assembly (M337V ALS-associated mutation and the helix-disrupting M337P engineered variant) both discourage helix-helix assembly and phase separation^32^. Given that the helix-stabilizing variants enhance interactions and phase separation, reducing backbone conformational entropy via substitutions at glycine residues in TDP-43 CTD conserved region appears to stabilize protein interactions at this site.

Importantly, the TDP-43 helical region appears unique among RNA-binding proteins – homologous (i.e. evolutionarily related) hydrophobic-residue-rich sequences are apparently not found in other human proteins and seem out of place amidst the polar (FUS, hnRNPA2, remainder of TDP-43 CTD) or highly charged (ddx4)^3^ low complexity sequences known to mediate LLPS. Yet, other phase separating proteins such as TIA-1 contain predominantly disordered domains with conserved helix-inducing residues (especially alanine which is enriched in the TDP-43 conserved region), suggesting that TDP-43 CTD may be one in a class of natural proteins containing helical modules that form upon and contribute to phase separation. Importantly, unlike traditional coiled coil or leucine zipper helix-helix interactions^50^, TDP-43 CTD conserved region encodes weak (*K*_d_ ~100 μM range) and dynamical helical interactions. This range of affinity is consistent with the multivalent but individually weak contacts formed within membraneless organelles and their dynamic, liquid nature where a tight contact (nM range) with slow off-rate could inhibit responsive reorganization of the functional complexes. Indeed, the apparent interaction strength is commensurate with self-assembly domains already characterized in the remainder of TDP-43: the polar-rich prion-like majority of the CTD that assembles via individually weak multivalent contacts32, and TDP-43 N-terminal domain that forms weak (*K*_d_ ~100 μM) globular protein chains also enhancing phase separation^26^. Similarly, the contacts formed by TDP-43 CTD conserved region are also heterogeneous^32^, unlike amphiphilic α-helices that form ordered, rigid bundles. Indeed, CTD assembly may underlie the ability of this region of TDP-43 to interact with splicing factor partners including hnRNPA2^31,43^. Hence TDP-43 CTD conserved region serves as a dynamic and heterogeneous binding module to enhance TDP-43 phase separation.

The identification of single, specific sites that can be used to substantially alter the assembly and material properties of cellular droplets offers an important opportunity to use TDP-43 CTD as a module to tune phase separation in engineered fusion proteins. The multivalent interactions leading to phase separation lend themselves to modulation^51^, and the TDP-43 CTD conserved region is shown here to be a uniquely short and tunable module. Interaction can be finely adjusted by single amino acid position changes to dramatically enhance or, as in our previous work, decrease interaction at this site (Figure 3). Enhanced interactions of this region in turn increases phase separation both *in vitro* (Figure 4) and in cells (Figure 5), leading to enhanced function in cells. Therefore, TDP-43 CTD helical region serves as a small structural element that can be used to perturb or control phase separation. Furthermore, compared with repetitive low complexity sequences (e.g. prion-like regions or arginine rich regions)^11^ or globular protein/ linear interaction motif pairs (e.g. SH3:proline-rich)^52^, TDP-43 CTD’s short (~20 residue) module functions with a fine ability to precisely tune phase separation to the desired degree, using the “knob” of mutations that can precisely increase or decrease interaction. Indeed, just as splicing can be altered by G335A that enhances helix-helix assembly of native TDP-43, variants of the TDP-43 conserved region could be fused to other proteins to regulate their cellular functions within membraneless organelles. Furthermore, just as phase separation of low complexity regions can be controlled by post-translational modification^46^, targeting modifications to sites adjacent to or within a TDP-43 helical module could be used to alter phase separation and function responsively. Therefore, a series of TDP-43 CTD variants could make for an important tool for engineering phase separation or, more broadly, for tuning dynamic self-organization.

## Acknowledgements

We thank Mandar Naik for assistance with NMR spectroscopy and Ashley Reeb for help with tissue culture and transfection experiments. Research was supported in part by NIGMS R01GM118530 (to N.L.F.), NSF 1845734 (to N.L.F.), and a starter grant 17-IIP-342 from the ALS Association (to N.L.F.). A.E.C. and A.M.D. were supported in part by NIGMS training grant to the MCB graduate program at Brown University (T32GM007601). G.L.D., G.Z., and J.M. are supported by the U.S. Department of Energy (DOE), Office of Science, Basic Energy Sciences (BES), Division of Material Sciences and Engineering under Award DESC0013979 (to J.M.). Work at Stanford was supported by grants from the NIH (DP2GM105448 and R35GM118082) to RR and a fellowship from the Deutsche Forschungsgemeinschaft (SCHM 3082/2-1) to HBS. This research is based in part on data obtained at the Brown University Structural Biology Core Facility supported by the Division of Biology and Medicine, Brown University. Use of the high-performance computing capabilities of the Extreme Science and Engineering Discovery Environment (XSEDE), which is supported by the NSF grant TG-MCB-120014, is gratefully acknowledged in addition to resources of the National Energy Research Scientific Computing Center, a DOE Office of Science User Facility supported by the Office of Science of the U.S. Department of Energy under contract DE-AC02-05CH11231. The content is solely the responsibility of the authors and does not necessarily represent the official views of the funding agencies.

## Methods

### Protein Expression and Purification

Point mutations were introduced into the TDP-43_267-414_ (CTD) construct previously described^32^ using Quikchange mutagenesis or overlap extension PCR with mutagenic primers^53^.

WT and mutant CTD constructs were expressed in *Escherichia coli* BL21 Star DE3 (Invitrogen) cells in either LB media or, where indicated as uniformly labeled ^15^N or ^15^N /^13^C, in M9 minimal media supplemented with ^15^NH_4_Cl and/or ^13^C -glucose as the sole nitrogen and carbon sources, respectively. Recombinant proteins were purified as previously described^32^, with the exception that the TEV cleavage buffer was substituted with 20 mM Tris, 500 mM GndHCl, pH 8.0 for more efficient cleavage of the purification tag. Additionally, proteins were exchanged into Storage Buffer (20 mM Tris, 8 M urea, pH 8.0) before flash freezing and storage at −80 °C. For all experiments, purified proteins were desalted into MES Buffer (20 mM MES, pH 6.1) using 0.5 mL Zeba spin desalting columns (Thermo Scientific) in accordance with manufacturer instructions.

### NMR Spectroscopy

All NMR experiments were preformed in 20 mM MES buffer pH 6.1 (pH adjusted with NaOH) supplemented with 10% D_2_O. Experiments were recorded on Bruker Avance spectrometers operating at either 850 MHz or 500 MHz ^1^H Larmor frequency with TCI z-gradient cryoprobes at 25° C. Backbone resonance assignments for CTD variants were obtained by standard triple resonance experiments HNCACB and CBCACONH as previously described^32^. Subsequent secondary chemical shifts were obtained from comparison of experimental C^α^ and C^β^ chemical shifts with their respective predicted random coil reference chemical shifts^54,55^.

^15^N *R*_1_, ^15^N *R*_2_, and {^1^H} ^15^N heteronuclear NOE experiments were measured for 20 μM WT, 20 μM G335A, and 10 μM - 20 μM G338A ^15^N-labeled CTD at 500 MHz ^1^H Larmor frequency using standard pulse sequences (hsqct1etf3gpsitc3d, hsqct2etf3gpsitc3d, hsqc-noef3gpsi). Each ^15^N *R*_1_ experiment comprises seven interleaved ^15^N *R*_1_ relaxation delays: 100 ms, 200 ms, 300 ms, 400 ms, 600 ms, 800 ms, and 1000 ms with a recycle delay of 1.5 s. Each ^15^N *R*_2_ experiment comprises six interleaved ^15^N *R*_2_ relaxation delays with a *B*_1_ field of 556 Hz: 16.4 ms, 32.8 ms, 65.5 ms, 131.2 ms, 229.6 ms, and 262.4 ms with a recycle delay of 2 s. {^1^H}-^15^N heteronuclear NOE experiments comprise interleaved proton saturation sequences and reference sequences without proton saturation with a recycle delay of 5 s. Measurement of *η*_xy_ was accomplished using the pulse sequence described previously^56^, with a relaxation delay (T) of 42 ms and recycle delay of 1.5 s.

Chemical shift perturbations were quantified for either WT and mutant CTD variants by measuring the chemical shifts of ^1^H-^15^N crosspeaks of HSQCs acquired at 850 MHz ^1^Η Larmor frequency at concentrations ranging from 10 μM to 90 μM as previously described^32^.

Pulsed-field gradient diffusion measurements for WT and mutant CTD at concentrations ranging from 10 μM to 600 μM acquired using the longitudinal eddy current delay with bipolar gradient scheme^57^ provided in the Bruker standard library (ledbpgppr2), with 16 gradient strengths: 0.96 G/cm, 5.39 G/cm, 9.92 G/cm, 13.62 G/cm, 16.58 G/cm, 20.10 G/cm, 23.75 G/cm, 27.26 G/cm, 30.83 G/cm, 34.37 G/cm, 37.90 G/cm, 41.38 G/cm, 44.82 G/cm, 48.08 G/cm, 51.33 G/cm, and 54.46 G/cm. The diffusion time was set to 250 ms with encoding/decoding gradient pulse lengths of 1 ms, using a recycle delay of 2 s. Gradient powers were calibrated based on the known diffusion coefficient of trace H_2_O in a 99.8 % D_2_O sample^58^. In all cases, intensity ratios were calculated by least-squares regression of select 1H spectral regions for each gradient power against those of a reference gradient power (0.96 G/cm). Standard error of the intensity ratios was estimated with 100 boostrapping simulations.

All NMR spectra were processed with NMRPipe^59^ and analyzed with NMRFAM-Sparky^60^. ^15^N *R*_1_, ^15^N *R*_2_, {^1^H} ^15^N NOE, and *η*_xy_ values and their respective standard errors were calculated using in-house scripts.

### Estimating *R*_ex_ for WT and G335A CTD

Chemical exchange contributions to ^15^N *R*_2_ (*R*_ex_) were calculated based on the relationship:

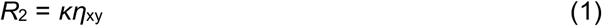

where *κ* is a scaling parameter that can be approximated for all residues as the trimmed mean of 61 *R*_2_*/η*_xy_^61^. In our case, two values of *κ* were calculated due to the differential relaxation rates across the CTD resulting from both structured and disordered regions. *η*_xy_ rates were first separated into one of two equal-width bins, and their respective *κ* values were obtained by solving equation (1). Site-specific *R*_ex_ values were then calculated from *R*_2_, *κ*, and *η*_xy_, as:

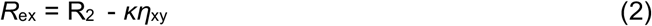

### Crosslinking

CTD variants harboring engineered cysteine residues at one of two serine positions (S273C or S387C) were buffer exchanged and diluted to 100 μM in Storage Buffer (see above, which includes 8M urea to maintain solubility). Crosslinking was initiated by addition of copper (II) phenanthroline^62^ to a final concentration of 1 mM, followed by incubation at room temperature for at least 1 hour. Reactions were quenched by addition of EDTA to 5 mM, and the resulting mixture was purified by size-exclusion chromatography with a 26/600 Superdex 75 equilibrated in Storage Buffer supplemented with 1 mM EDTA. Fractions corresponding to dimeric CTD were pooled, concentrated up to 750 μM ([dimer]), flash frozen, and stored at −80C. NMR samples were prepared by diluting 12.5 μL of the dimeric stock into 437.5 μL MES Buffer, resulting in a final urea concentration of 200 mM. Samples were then centrifuged for 10 minutes at 12,000 rpm, and the concentration of the supernatant was measured by A_280_. The samples were then diluted to the final desired protein concentration with 1:35 Storage Buffer:MES Buffer, thus ensuring the urea concentration remains constant. In order to prepare monomeric control samples, the samples were supplemented with 1 mM DTT (Sigma) prior to centrifugation and at all subsequent steps.

### Quantifying phase separation *in vitro*

WT and mutant CTD variants were desalted into MES Buffer and diluted to 40 μM. Phase separation was initiated by adding an equal volume of MES Buffer supplemented with either 0, 75, 150, 300, 600, or 1000 mM NaCl, resulting in final salt concentrations of 0, 37.5, 75, 150, 300, and 500 mM NaCl, respectively. Samples were mixed by gentle agitation and promptly centrifuged at 12,000 rpm for 10 min. The concentration of CTD in the supernatant was measured by A_280_ (*E*_280_ = 17990 M^−1^cm^−1^) with a Nanodrop 2000c. All experiments were performed in triplicate.

### Microscopy

WT and mutant CTD were desalted into MES Buffer and diluted to 40 μM. Phase separation was initiated by addition of an equal volume of MES Buffer supplemented with either 0 mM or 600 mM NaCl, resulting in final salt concentrations of 0 or 300 mM NaCl, respectively. Samples were spotted onto glass coverslips, and differential interference contrast (DIC) micrographs were acquired with a Zeiss Axiovert 200M as previously described^32^. Images were processed with ImageJ.

### Translational Diffusion

Translational diffusion coefficients of WT and mutant CTD were calculated by non-linear least squares fit of CH_3_ ^1^H spectral region intensity ratios to the Stejskal-Tanner equation^63^:

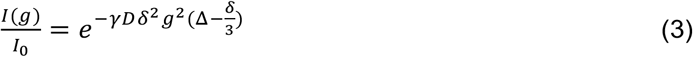

where *γ* is the ^1^H gyromagnetic ratio, *D* is the translational diffusion coefficient, *δ* is the length of the bipolar encoding/decoding gradient element, Δ is the diffusion time, and *g* is the gradient strength. Standard error in the diffusion coefficients was estimated from the variance-covariance matrix of nonlinear least-squares model parameters.

### TDP-43_RRM-GFP_ reporter assays

HEK-293T cells (ATCC catalog number CRL-3216; RRID:CVCL_0063) maintained in DMEM high glucose (GE Healthcare) supplemented with 10% FBS, 2mM L-glutamine (Gibco), 1mM non-essential amino acids (Gibco) and 1% penicillin/streptomycin (Gibco) at 37°C and 5% CO_2_ were seeded onto ibidi μ-slides at 30,00 cells/well and transfected with TDP-43_RRM-GFP_ reporter constructs (300 ng) using X-tremeGENE 9 (Roche). After 24 hours, the culture medium was exchanged with L-15 medium (Gibco) supplemented with 10% FBS, and the cells were imaged using a Leica SP8 laser-scanning confocal microscope equipped with a 488-nm laser, 63x glycerol objective (NA 1.3) and a temperature-controlled incubation chamber (Life Imaging Services) set to 37°C.

Half-bleach experiments were recorded and quantified using the FRAP module of the Leica Application Suite X software as previously described^33^. In short, for each recorded time point (t), the fluorescence intensities within the bleached droplet hemisphere were then normalized to the fluorescence intensity of the corresponding unbleached droplet hemisphere. These normalized, time-dependent fluorescence intensities It were then used to calculate the fluorescence recovery (FR) according to the following formula: FR(t) = (I_t_ − I_t0_) / (I_before bleaching_ − I_t0_), with t0 being the first time point observed after photobleaching. Two replicate experiments per construct were performed and at least ten individual droplets from different cells were analyzed per experiment. Replicate measurements (mean ± SD) were plotted using GraphPad Prism. To ensure that only similarly-sized droplets of comparable intensities were bleached, the same bleach window and detector settings were used. Droplet diameters and fluorescence intensities were quantified using Mathematica (Wolfram Research) as previously described^33^.

### Splicing assays

WT and mutant siRNA-resistant constructs were generated by site-directed mutagenesis in N-terminally FLAG-tagged pCMV2^64^. HeLa cells were grown in growth media-Dulbecco’s modified Eagle’s medium, 4,500 mg/l glucose, l-glutamine, and sodium bicarbonate and supplemented with filtered fetal bovine serum at 10%. Cells were transiently transfected with Lipofectamine 2000 and Oligofectamine in the case of siRNA (Life Technologies) according to manufacturer protocols. RNAi-mediated downregulation of TDP-43 was carried out as previously described^64^ and siCONTROL Nontargetting siRNA#1 (Dharmacon) was used as control. TDP-43 downregulation was confirmed by immunoblotting. Splicing assays were performed using the CFTR reporter minigene as previously described^64^. WT or mutant siRNA-resistant vectors were co-transfected in T2-treated cells. To quantify exon skipping we performed standard PCR with primers specific for exon 9 flanking regions following RNA extraction and cDNA synthesis. The percent exon exclusion was calculated using ImageJ analysis.

### All-atom simulations

Parallel tempering in the well-tempered ensemble (PT-WTE) simulations^65,66^ were conducted on a 41-residue fragment of TDP-43 CTD, TDP-43_310*–*350_ using the GROMACS-4.6.7 molecular dynamics software package^67^. We use the Amber03ws force field, which has been developed and parameterized for use with disordered proteins and protein folding^68^. For simulations of WT TDP-43_310-350_ dimer, we also apply a well-tempered metadynamics (WT-MetaD) bias^69^ on two collective variables, intermolecular hydrophobic contacts and intermolecular hydrogen bonds to aid in sampling bound and unbound states.

### Coarse-grained simulations

Coarse-grained simulations were conducted using an amino-acid-resolution model with 20 residue types to capture sequence specificity, having interactions based on relative hydropathies of each amino acid^48,70^. Each system was simulated at a range of temperatures using constant volume and temperature using a Langevin thermostat, and slab geometry, which allows for efficient sampling of coexistence densities. Simulations of phase coexistence were conducted using HOOMD-Blue v2.1.5 software package^71,72^. For simulations of helical TDP-43, residues 320-334 were constrained to a helical configuration using rigid body dynamics.

### Analysis of Simulations

Secondary structures were calculated using the DSSP algorithm^73^, and secondary structure maps are used to show heterogeneity of helicity within the equilibrium ensemble^39^. From the PT-WTE-MetaD dimer simulation, we generate the free energy profile with respect to the two collective variables^74^. All analysis done on the dimer simulation was reweighted based on the applied bias. Chemical shifts were calculated using SPARTA+^75^, and normalized using random-coil shifts from the Poulson webserver^55^.

## Data Availability

NMR chemical shift assignments in this paper for TDP-43 CTD G335A (27750), G335N (27788), G335S (27789), G338A (27790), and G335D (in process) can be obtained online from the Biological Magnetic Resonance Database (BMRB, http://www.bmrb.wisc.edu/). TDP-43 CTD WT chemical shifts were obtained from the BMRB (26823). Plasmids generated for this project can be found at Addgene.org.

## Author Contributions

JM and NLF conceived and designed research. AEC and AMD conducted biophysical experiments. GLD and GHZ performed and analyzed simulations. HBS and YMA conducted in-cell experiments. YCK developed tools to aid in setup and analysis of simulations. RR, YMA, JM, and NLF funded and supervised research. AEC, GLD, JM, and NLF wrote manuscript.

---

**SI Figure 1:**
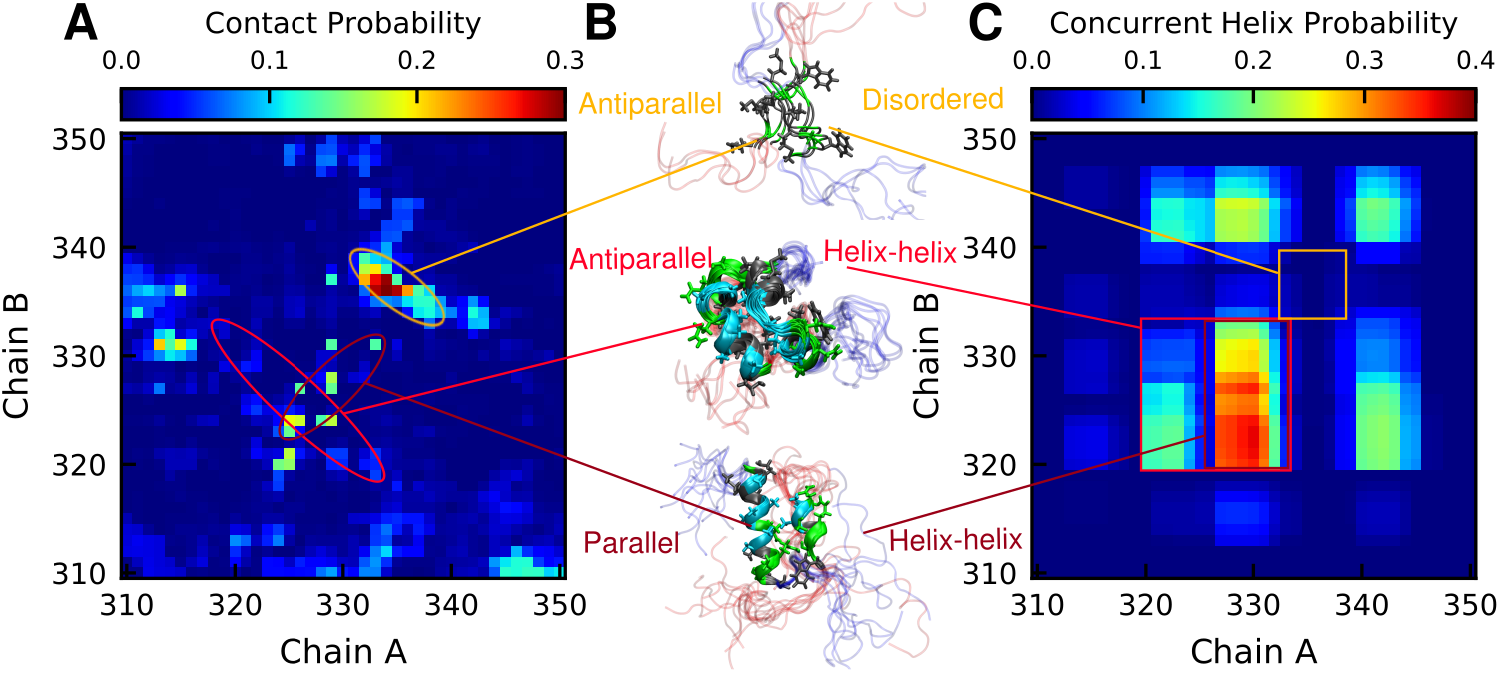
Self-association of TDP-43 CTD vs helical content. A) Intermolecular contact probability map of two chain simulation of TDP-43_310-350_ calculated by distance between *α*-carbon atoms of each residue. Cutoff value for contacts is set to 8 Å. B) Clusters of bound configurations highlighting regions that are in contact, and helical with the exception of the disordered contact on the top snapshot. C) Per-residue probability of concurrent helix formation between the two chains, showing which regions of the protein are commonly helical at the same time.

**SI Figure 2:**
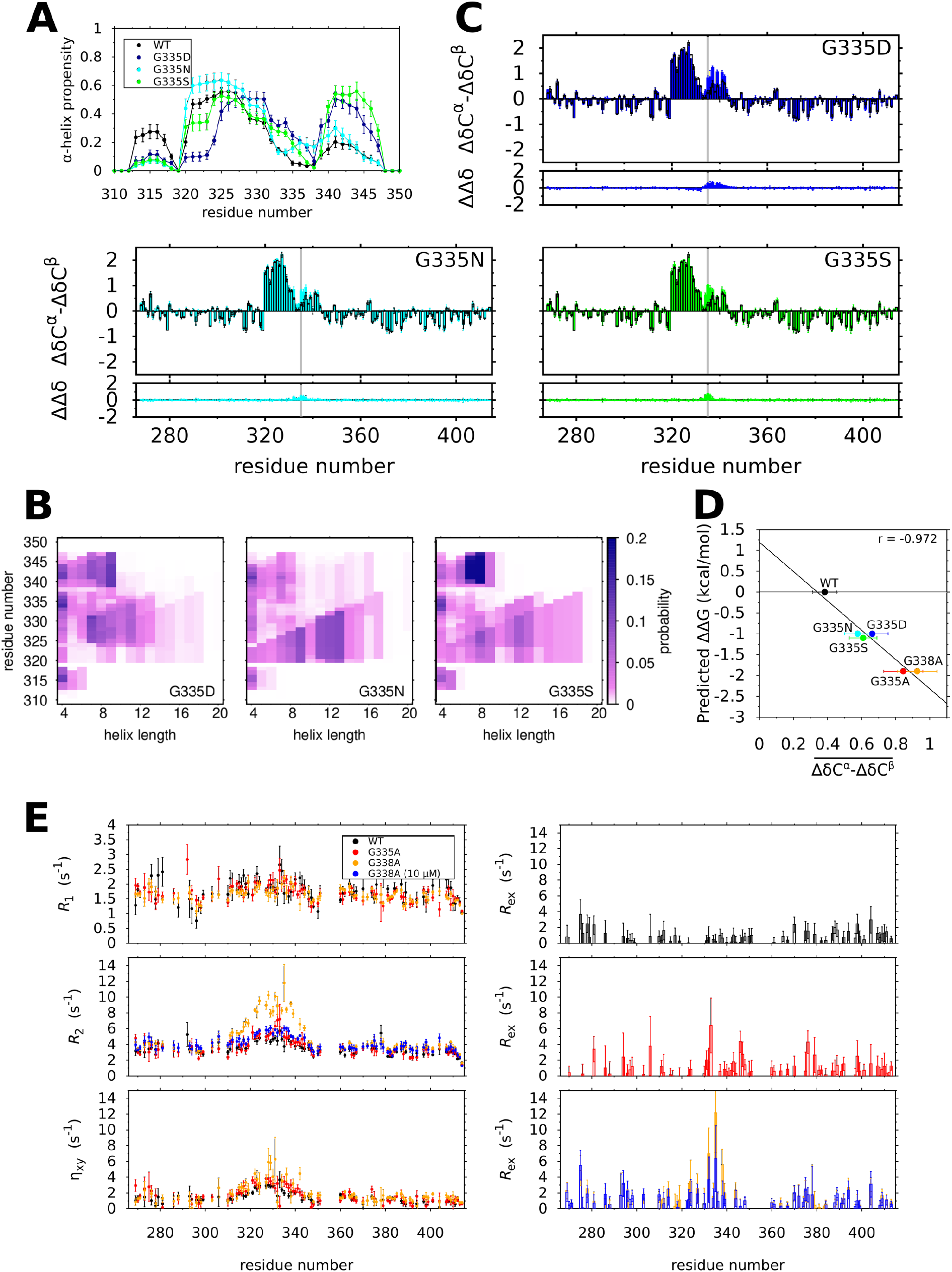
Mutations at G335 and G338 enhance local *α*-helical structure. A) *α*-helical propensities for WT, G335D, G335N, and G335S CTD_310-350_ determined by all-atom MD simulation. An unexpected dip for G335D far from the site of mutation (residues 320-324) may be due to a change in net charge of the peptide. Error bars are SEM. B) Secondary chemical shifts (**ΔδC**^***α***^ − **ΔδC**^***β***^) and the differences in secondary shifts with respect to WT (ΔΔδ) for G335D, G335N, and G335S CTD. Error bars are SD. C) Predicted change in free energy ΔΔG for monomeric *α*-helix formation versus the average change in secondary shift 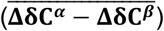 for residues 331-343 of G335 and G338 variants. D) ss-maps for G335D, G335N and G335S TDP-43_310-350_ E) ^15^N spin-relaxation parameters *R*_1_, *R*_2_, and η_xy_, as well as calculated *R*_ex_ values are plotted for WT, G335A, and G338A at 10-20 μM sample concentration. Error bars are SD.

**SI Figure 3:**
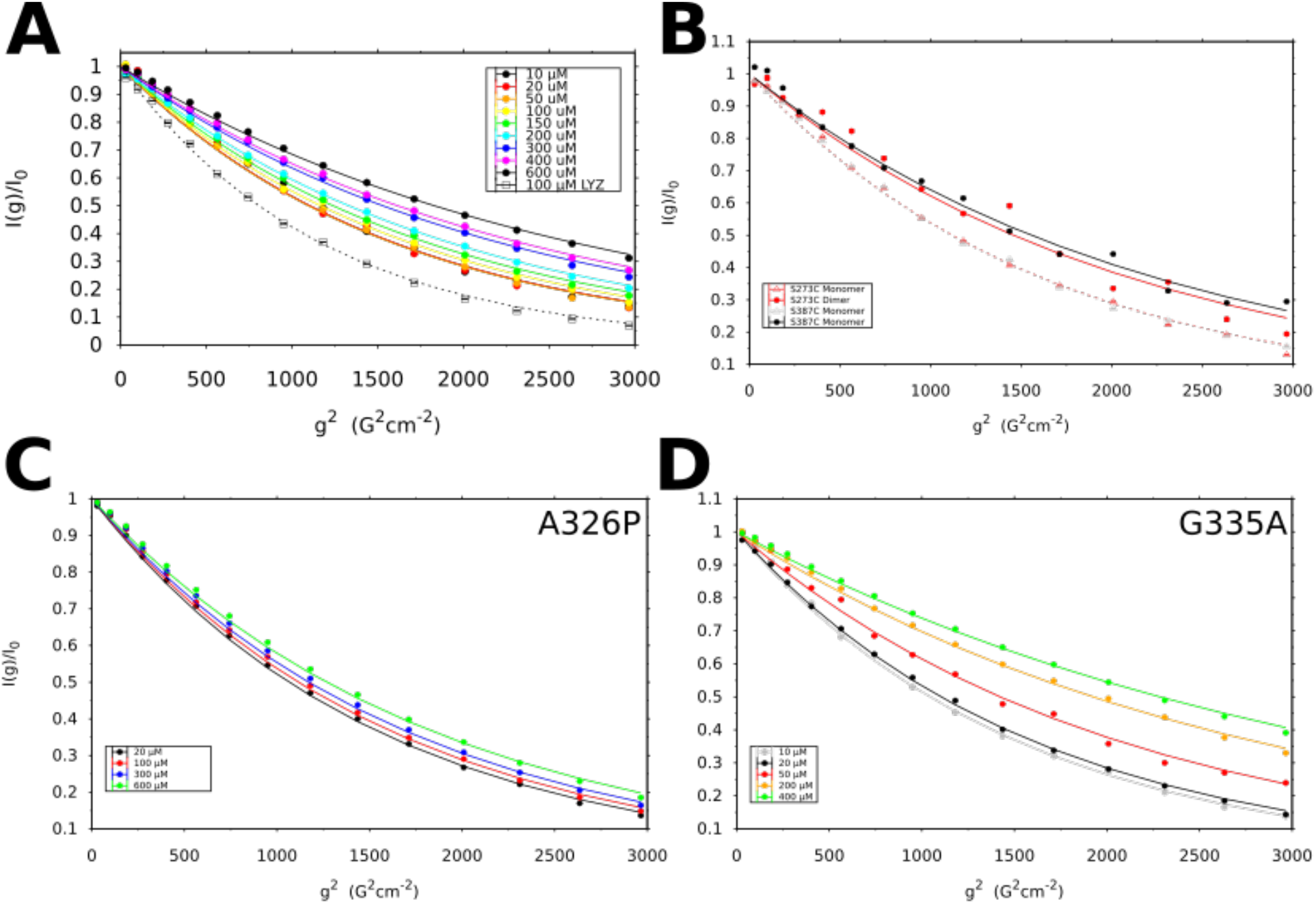
Mutations at G335 and G338 enhance intermolecular helix-helix contacts and higher-order assembly. A) Diffusion NMR displayed as normalized resonance intensity (I(g)/I_0_) versus squared gradient strength (g^2^) show that diffusion rate changes rapidly for WT CTD from 150 μM to 300 μM. Lines are best fit to the Stejskal-Tanner equation. Control sample of 100 μM lysozyme (LYZ) shown as black dashed line. Error bars are SD from 100 bootstrapping simulations. Intensity ratios of methyl ^1^H resonances for monomeric/dimeric S273C and S387C B), A326P C), and G335A D) plotted as a function of squared gradient strength (g^2^). Error bars are SD from 100 bootstrapping simulations. The solid and dashed lines represent the best-fit nonlinear-least squares solution of the Stejskal-Tanner equation.

**SI Figure 4:**
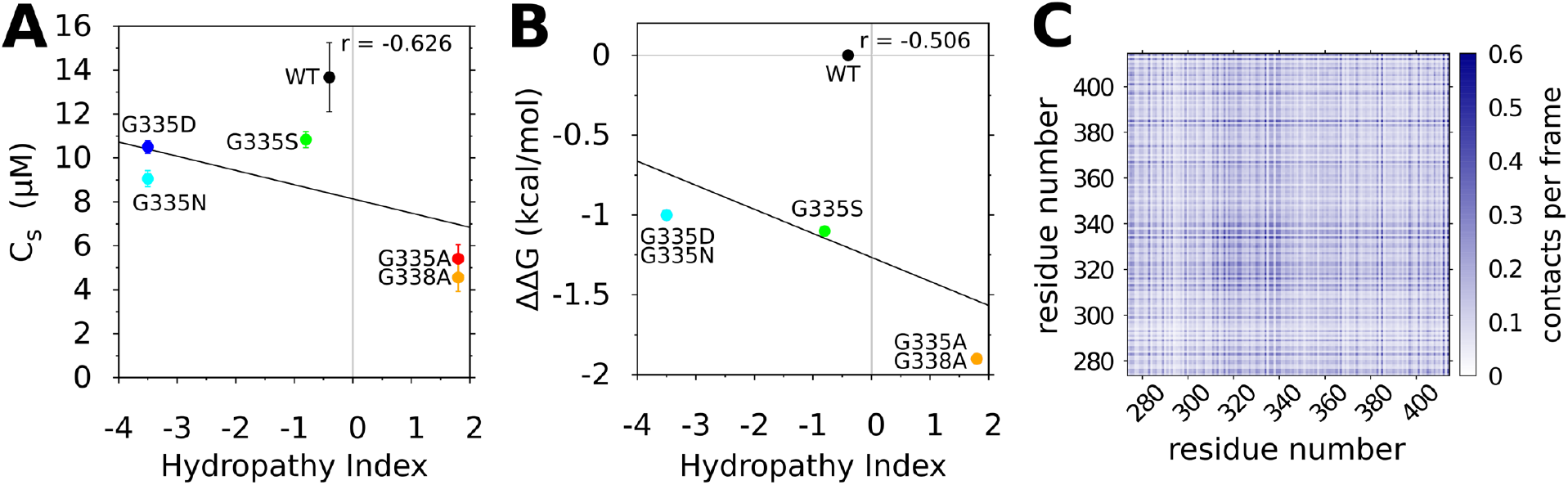
Mutations at G335 and G338 alter *in vitro* LLPS. (A) Correlation plot of C_s_ values at 150 mM NaCl versus hydropathy index based on the scales described by Kyte and Doolittle^76^ shows poor correlation, suggesting that changes in residue hydrophobicity of G335 and G338 variants do not explain changes in phase separation. (B) Correlation plot of the change in free energy formation for a given amino acid to adopt monomeric α-helical structure with respect to glycine (ΔΔG) versus hydropathy index. C) Intermolecular contact map for coarse-grained slab simulation of TDP-43 CTD highlights residues 314-340 as having the highest propensity of intermolecular interaction in the equilibrated simulation ensemble.

**SI Figure 5:**
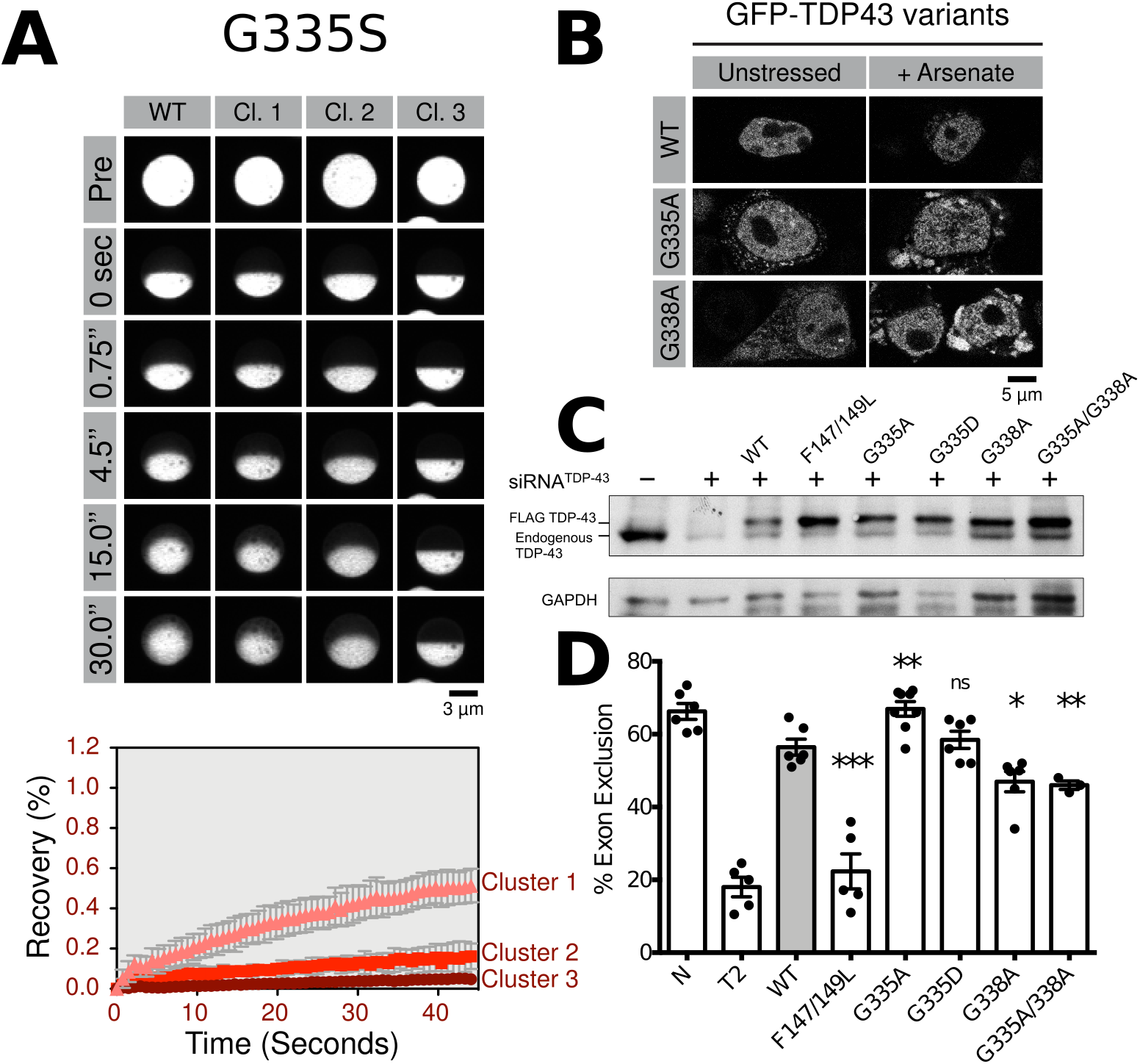
Mutations at G335 and G338 alter in-cell LLPS and splicing function. A) Fluorescence recovery after in cell half-droplet photobleaching for G335S TDP-43_RRM-GFP_ illustrates heterogeneity of G335S droplet fluidity. Observations of TDP-43 reporter droplet fluidities from individual cells were manually grouped by FRAP recovery rates into clusters to demonstrate the heterogeneity that gives rise to large uncertainties for G335S data analyzed without grouping (seen for G335S in Figure 5B). Scale bar is 3 μm. B) Fluorescence microscopy of GFP-TDP-43 WT, G335A, and G338A highlight the formation of cytosolic puncta in mutant variants after arsenate-induced cellular stress. Scale bar is 5 μm. C) Western blot of G335 and G338 variants used in exogenous CFTR exon 9 minigene assay shows protein expression levels between WT and mutant variants that are comparable when compared to GAPDH loading control variations. WT and mutant constructs were expressed in TDP-43 siRNA-treated HeLa cells. D) Percent CFTR exon 9 skipping measured from control and siRNA TDP-43 (T2) treated HeLa, expressing siRNA-resistant WT and mutant constructs. Non-specific siRNA was used as control (N). Statistical significance and SEM calculated for n≥3, using unpaired, two-tailed, Welch-corrected t-test (P *** P = 0.0008, ** P = 0.004, and *P = 0.02).

